# Understanding the Evolution of Nutritive Taste in Animals: Insights from Biological Stoichiometry and Nutritional Geometry

**DOI:** 10.1101/2021.03.11.434999

**Authors:** Lee M. Demi, Brad W. Taylor, Benjamin J. Reading, Michael G. Tordoff, Robert R. Dunn

**Affiliations:** Department of Applied Ecology, North Carolina State University, Raleigh, NC 27695, USA; Monell Chemical Senses Center, Philadelphia, PA 19104, USA; Center for Evolutionary Hologenomics, University of Copenhagen, Copenhagen, DK

**Keywords:** gustation, nutritional ecology, chemoreception, homeostasis, optimal foraging

## Abstract

A major conceptual gap in taste biology is the lack of a general framework for understanding the evolution of different taste modalities among animal species. We turn to two complementary nutritional frameworks, biological stoichiometry theory and nutritional geometry, to develop hypotheses for the evolution of different taste modalities in animals. We describe how the attractive tastes of Na, Ca, P, N and C containing compounds are consistent with principles of both frameworks based on their shared focus on nutritional imbalances and consumer homeostasis. Specifically, we suggest that the evolution of multiple nutritive taste modalities can be predicted by identifying individual elements that are typically more concentrated in the tissues of animals than plants. Additionally, we discuss how consumer homeostasis can inform our understanding of why some taste compounds (i.e., Na, Ca and P salts) can be either attractive or aversive depending on concentration. We also discuss how these complementary frameworks can help to explain the phylogenetic distribution of different taste modalities and improve our understanding of the mechanisms that lead to loss of taste capabilities in some animal lineages. The ideas presented here will stimulate research that bridges the fields of evolutionary biology, sensory biology and ecology.

## 1. Introducing a general framework for consumptive taste evolution

All organisms are faced with the challenge of procuring the essential elements and biochemicals of life for growth, maintenance and reproduction, often in proportions that deviate substantially from their environmental availability. For animals, nutrients (i.e., elements and/or biochemicals) are acquired through their diet by consuming foods that may vary considerably in their chemical composition and that frequently provide an inadequate supply of one or more essential nutrient (1). Thus, many animals suffer periodic nutritional imbalances whereby the nutrient content of available foods does not match their nutritional requirements. Such imbalances (i.e., over- or undersupply of essential nutrients) can affect consumer metabolism and performance, ultimately resulting in reduced growth and fitness (2, 3). Because nutritional imbalances between consumers and their foods are both pervasive and consequential animals have evolved a range of adaptations, from behavioral to physiological and chemosensory, that allow them to modulate nutrient intake, as well as post-ingestion nutrient processing (i.e., assimilation and allocation), to minimize potential imbalances (3).

Selective foraging is a key behavioral adaptation that allows animals to regulate nutrient intake in order to achieve balanced nutrient supply (3). Choosing exactly which foods to eat, however, requires that animals differentiate among potential food sources of differing chemical composition to select those that are most nutritionally advantageous. One of the primary mechanisms that animals use to assess the nutritional quality of foods is gustation, or taste, which provides animals the ability to evaluate the chemical composition of potential foods prior to ingestion via complex chemosensory systems (4, 5, 6). Indeed, there is broad recognition that gustatory systems function primarily as a screening mechanism that drives consumption of nutrient- and energy-dense foods, as well as the rejection of potentially harmful ones based upon the presence, and concentrations, of a variety of different chemicals. These various chemical stimuli interact with specialized receptor cells to produce signals that are transduced and interpreted as unique taste modalities, including the salty, sweet, bitter, sour and umami tastes familiar to humans (4, 6).

The variety of different taste modalities provide organisms with a way to evaluate the quality of food based on multiple aspects of its chemical composition and can be broadly grouped by whether they taste “good”, and therefore promote consumption, or “bad”, thus producing an aversive response. Consumptive responses, such as those elicited by sweet and umami tastes, promote the ingestion of foods that supply compounds essential for growth and metabolism. Conversely, aversive responses, such as those typically generated by bitter and sour tastes, encourage animals to reject, rather than consume specific foods. Aversive tastes are generally thought to inform consumers that food may contain toxic or harmful chemicals, such as allelochemicals, though not all chemicals that produce innate aversive responses are necessarily harmful (5, 6). Additionally, some elements, such as Na, Ca and P which are essential but can be toxic when oversupplied, may elicit either consumptive or aversive responses depending on concentration, thereby allowing consumers to regulate intake within a relatively narrow window.

All animals possess gustatory sensing capabilities (3, 7), reinforcing the notion that regulation of nutrient intake is highly consequential for consumer fitness. Interestingly, gustatory sensing appears to have evolved largely independently among prominent animal lineages (i.e., mammals and insects) (5), yet exhibits remarkable convergence around taste capabilities for a small suite of compounds (e.g., amino acids, carbohydrates, Na salts). This suggests that animals have faced similar nutritional constraints throughout their evolutionary history and indicates potential for developing a nutritional framework for understanding the evolution of taste that has broad applicability. Nevertheless, we currently lack a predictive understanding of when and why particular taste modalities might be evolutionarily favored (3), thereby leading to differences in the breadth of gustatory capabilities among species.

In this paper, we turn to two complementary frameworks, biological stoichiometry theory (2, 8, 9) and nutritional geometry (1, 3, 10), that have been developed by ecologists to characterize nutritional imbalances (and their consequences) in trophic interactions to draw insights into the evolutionary contexts under which different taste modalities have evolved in animals. In the following sections, we summarize the basic tenets of biological stoichiometry theory and nutritional geometry, including the unifying concept of consumer homeostasis, and draw upon a review of the taste biology literature to show how these complementary frameworks advance our understanding of the evolution of multiple taste modalities in animals. We use the principles of biological stoichiometry to demonstrate that the evolution of multiple nutritive taste modalities can be predicted by identifying individual elements that are typically more concentrated in the tissues of animals than plants. Moreover, we describe how consumer homeostasis can inform our understanding of why some taste compounds (i.e., Na, Ca and P salts) can be either attractive or aversive depending on concentration. Additionally, we demonstrate how these complementary frameworks can help to explain the phylogenetic distribution of different taste modalities, including why the taste of Ca (at least at some concentrations) is attractive to some vertebrate species but appears to be strictly aversive to insects. We also discuss how biological stoichiometry and nutritional geometry can improve our understanding of the mechanisms that lead to loss of taste capabilities, as well as the pseudogenization of associated taste receptor genes, in some animal lineages (11, 12, 13). We also discuss some caveats to our application of these complementary frameworks and finally, propose several ideas that we hope will stimulate future research, including into the evolutionary history and phylogenetic distribution of different taste modalities, and incorporation of taste into functional trait-based ecological analyses.

## Nutritional frameworks in ecology

Ecologists have long recognized that the quantity and nutritional quality of foods affect the growth and performance of animals (1). Given that nutritional requirements vary among animal species, shifts in the nutrient composition of available foods can scale beyond individual and population effects to alter community and ecosystem dynamics, including fluxes of energy and nutrients to and from the environment and among trophic levels (14, 15). Increasing recognition of the role of animal nutrition in driving ecological dynamics, from foraging behavior to consumer driven nutrient recycling, has led to the development of multiple conceptual frameworks that seek to characterize the nature and consequences of nutritional imbalances in trophic interactions (1). Biological stoichiometry theory and nutritional geometry are perhaps the most prominent of these frameworks and have been developed in parallel over the last several decades (14, 15). Though these frameworks focus on different nutritional currencies (i.e., elements versus macro- and micronutrients) there is substantial conceptual overlap, including an emphasis on homeostasis, that suggest promise in their ability to inform our understanding of how nutritional imbalances have shaped the evolutionary history of gustation in animals (14). There are large bodies of literature devoted to the development and application of these frameworks, including several recent papers highlighting their connections and encouraging their synthesis (14, 15, 16). As such, we provide only brief summaries of their basic principles to establish their usefulness in describing the ecological contexts under which different taste modalities have evolved.

Biological, or commonly, ecological stoichiometry is the study of the balance of energy and multiple elements in ecological interactions (2, 8, 9). For the purposes of this paper, we prefer the term “biological stoichiometry”, as its broader scope explicitly integrates cellular and genetic mechanisms (9). Similar to stoichiometry of chemical reactions, biological stoichiometry applies thermodynamic principles to the mass balance of multiple elements in biological interactions (8). It is rooted in the observation that individual organisms are composed of elements in relatively fixed ratios (stoichiometric homeostasis) and that available resources frequently supply elements in ratios that deviate from those of the organism. This is particularly true of animals, which tend to be far less plastic in their elemental composition and exhibit far narrower interspecific variation in elemental composition than plants (and many autotrophic microorganisms; Fig. 1), thereby resulting in notable stoichiometric imbalances between plants and animals (17; 18). Core principles of biological stoichiometry include the observations that (1) organisms have limited capacity to alter their elemental composition even when the supply of elements is variable and that (2) large imbalances in elemental composition between consumers and their foods (e.g., herbivores and plants, Fig. 1) are common (2). A central thesis of biological stoichiometry theory is that reconciling these imbalances (i.e., via selective foraging behavior or post-ingestive regulation) while maintaining elemental homeostasis has consequences for the growth and fitness of animals, as well as the cycling of elements between animals and the environment. Hence, a key strength of biological stoichiometry is the ability to scale and make predictions across different levels of organization (e.g., from consumer taste receptors to ecosystem nutrient supply).

**Fig 1.**
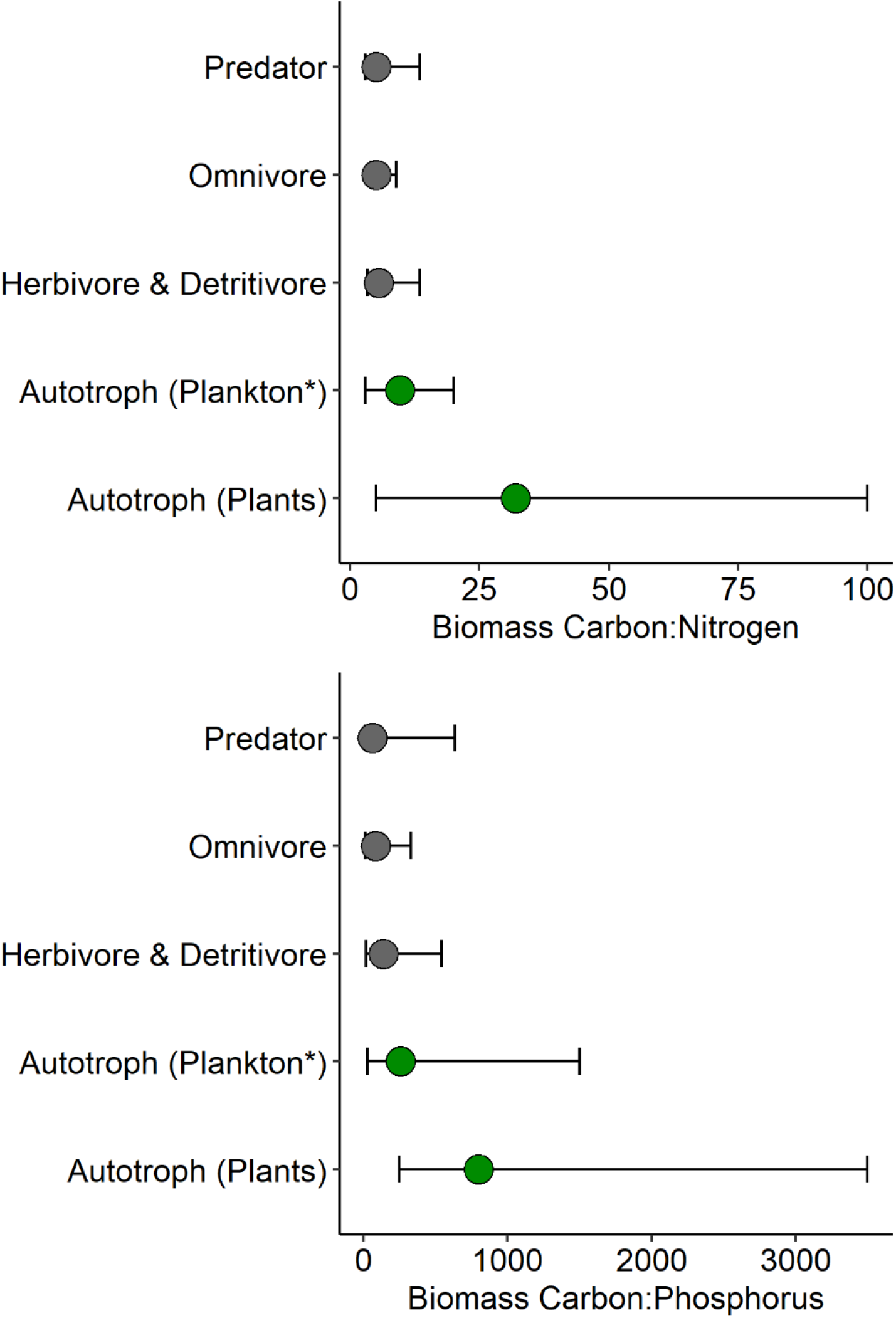
Carbon:nitrogen and carbon:phosphorus ratios of biomass across multiple trophic levels. Circles represent median values, error bars indicate ranges of C:N and C:P observed for organisms within each trophic level. Autotroph data are from Elser et al. (9), where Autotroph (Plants) includes measurements of foliar chemistry of terrestrial autotrophs (for C:N, n = 406; for C:P, n = 413) and Autotroph (Plankton*) represents nutrient chemistry of seston from freshwater lakes (for C:N, n = 267; for C:P, n = 273). Seston contains some detritus and heterotrophic biomass (i.e., bacteria, protozoa) but is typically dominated by phytoplankton (9). Consumer stoichiometry data are from Vanni et al. (79) and represent primarily aquatic animals. Data include 190 families from 9 animal phyla (Herbivore and Detritivore, n = 168; Omnivore, n = 77; Predator, n = 163).

Biological stoichiometry has historically focused primarily on imbalances between the supply and demand of C, N and P in trophic interactions. This bias reflects how commonly these elements limit biological productivity across trophic levels in many different ecosystems (19, 20) and a tendency to view limitation through the prism of single nutrient limitation rooted in Liebig’s law of the minimum. However, biological stoichiometry is not constrained to these three elements and incorporation of a wider suite of biologically essential elements into stoichiometric analysis is an emerging frontier (21, 22). Additionally, there is increasing recognition that key molecular and cellular level processes may be co-limited by multiple chemical elements (23) and that studies of nutrient limitation should consider this possibility and expand the elements considered in stoichiometric analyses (24). A notable limitation of biological stoichiometry theory with respect to its ability to characterize nutritional imbalances is its focus on individual elements (25, 14). Given that animal nutrition is largely biochemical in nature, focusing solely on the ratios of elements in foods may obscure potential imbalances in the supply of certain essential macro- and micronutrients that affect consumer performance and reduce fitness (25).

Similar to biological stoichiometry, nutritional geometry, or the “geometric framework” for animal nutrition, focuses on how the availability of multiple nutrients affect consumer growth and fitness and relies on homeostatic regulation as a conceptual cornerstone (1, 3, 26). However, nutritional geometry differs by focusing on the availability of micro- and macronutrients (i.e., amino acids, carbohydrates, lipids), rather than chemical elements, making it a more nutritionally explicit framework (1, 14). This framework recognizes that foods are complex mixtures of nutrients and other non-nutritive compounds and that any single food item is likely to provide an insufficient supply of at least some essential nutrients relative to consumer requirements at a given point in time. Nutritional geometry models relationships among various nutrients (or non-nutrients, such as allelochemicals in some applications) using a multi-dimensional state-space approach to characterize the mechanisms, such as foraging behavior or nutrient assimilation, that animals use to achieve nutritional homeostasis while foraging among foods of varying nutrient composition (26). This geometric approach was designed to provide quantitative predictions regarding how consumers utilize multiple foods that vary in their respective amounts of different nutrients (i.e., proteins and carbohydrates) in the service of homeostatic regulation of nutrient intake (1, 26).

Much of the early formulation and application of nutritional geometry focused on how animals forage among various resources to achieve a specific target ratio of two or more nutrients in their diet (14). Proponents of nutritional geometry have recognized that taste and other chemosensory systems (i.e., olfaction), as well as learned nutritional associations for various foods, play an important role in nutrient-based foraging decisions, allowing consumers to achieve balanced supply of multiple nutrients (3). In contrast, most applications of biological stoichiometry have focused on how organisms reconcile imbalances in the supply of essential elements via post-ingestive mechanisms, such as altering element use efficiencies (i.e., assimilation) and differential excretion of individual elements, rather than through selective foraging. However, the potential for biological stoichiometry theory to inform our understanding of how animals forage among nutritionally heterogenous food resources, particularly through the evolution and employment of chemosensory systems, has been previously suggested (17), but remains largely unexplored.

In the following sections we describe how these two complementary frameworks provide a platform, rooted in the concept of consumer homeostasis, to describe the evolutionary context from which multiple different taste modalities have evolved. We rely primarily on the more reductionist approach of biological stoichiometry theory, which focuses on nutritional imbalances at the elemental level, for many examples, including hypotheses related to evolutionary drivers of taste associations for N, P, Na and Ca containing molecules. We acknowledge that our use of biological stoichiometry at times treats individual elements as surrogates for complex biochemicals, and that gustatory receptors may be responding to those biochemicals rather than to individual elements contained within them, as in the case of N and amino acids. However, this approach remains useful in most cases given that a standard assumption of biological stoichiometry is that tissue concentrations of individual elements can often be related to a small number of specific biochemicals, such as protein being the primary N pool in most organisms (2, 14). There are, however, notable cases where more biochemically focused nutritional geometry provides greater insight into potential evolutionary drivers of taste, such as for carbohydrate and lipid taste. As such, we rely on both frameworks in our discussion of the evolutionary contexts for taste modalities, with the common thread being that consumers have relatively specific nutritional requirements, determined largely by their own chemical composition (which is relatively homeostatic), and that potential foods (at least individually) often do not satisfy those requirements.

## 3. Consumer homeostasis and nutritional imbalance: an evolutionary context for nutritive taste

The concept of elemental homeostasis, meaning that organisms have limited capacity to alter their chemical composition even when the supply of chemical elements is variable (2), is foundational to both biological stoichiometry theory and nutritional geometry. A second shared concept is that for many animals, the chemical composition of potential foods can be very different than that of the consumer. These differences result in nutritional imbalances, where certain foods may provide either an under- or oversupply of certain essential nutrients (i.e., elements or biochemicals), that can constrain individual performance and reduce reproductive potential (2, 3). Because consumers have limited capability to modify their chemical composition to more closely match that of their foods, they must reconcile imbalanced nutrient supply by using pre- and/or post-ingestive mechanisms (2, 3, 27), such as selective feeding or differential release of individual elements as waste products to achieve nutritional, or maintain tissue, homeostasis. However, post-ingestion nutrient processing, particularly of nutrients which are oversupplied, may have metabolic costs that affect consumer performance, highlighting the importance of selective foraging to reduce nutritional imbalances at the front end (28, 29). Gustation is commonly viewed as having evolved to function as a sensory tool that allows animals to assess the chemical composition of their food. Expanding further, we suggest that the evolution of many individual taste modalities should be explained, at least in part, by common nutritional imbalances in the diets of animals. Moreover, we contend that these common imbalances are the drivers of the remarkable convergence in gustatory evolution among major animal lineages (5). We believe that biological stoichiometry and nutritional geometry provide useful frameworks for identifying and characterizing those imbalances and therefore have great potential to inform our understanding of the evolutionary contexts for many individual taste modalities in animals.

Biological stoichiometry in particular provides a simple, yet useful, framework for characterizing common nutritional imbalances between consumers and their foods by comparing their respective elemental composition (though see carbon, section 4). We can make some useful generalizations about consumer nutritional imbalances by identifying elements that are commonly less concentrated in foods than in the tissues of consumers. The common convention in biological stoichiometry is to express the relative concentrations of different elements of consumers and their food as ratios, particularly as C:nutrient (i.e., N or P) ratios. For example, Fig. 1 displays biomass C:N and C:P ratios of autotrophs (i.e., plants) and animals across multiple trophic levels. The median C:N and C:P of autotroph biomass is considerably greater than that of animals (at any trophic level; Fig. 1), indicating that plant tissue has a greater amount of C per unit N, and C per unit P, than does animal tissue. As such, an animal of typical chemical composition (i.e., around the median) that is feeding on a plant of typical chemical composition will experience undersupply of both N and P, relative to C. We would expect that such elements, here N and P, would have a greater likelihood of limiting consumer growth and metabolism and therefore predict that animals should find them (or the primary chemicals that contain them, as in the case of N and amino acids) pleasing to taste - the pleasing oral sensation would reward animals for consuming foods that are relatively high in those potentially limiting nutrients.

To illustrate, we compared the whole-body elemental composition of insects, marine fishes, and mammals with the elemental composition of foliar plant tissue (Fig. 2, Table 1) in order to identify which elements are typically less concentrated in plants than in animals. We use plants and animals in this example because of the typically greater difference in the elemental composition of herbivores and plants than carnivores and their animal prey (Fig. 1) and with the assumption that this is representative of conditions faced by herbivorous animals (particularly terrestrial) throughout their evolutionary history. Based on principles of biological stoichiometry, we might expect there to be consumptive taste associations for elements that are typically more concentrated in the tissues of animals than in plants, given that these elements are likely to limit performance and reproductive potential. By extension, we would not necessarily expect positive taste associations for elements that are more concentrated in plants than animals. Comparison of animal and plant elemental composition reveals that Na, P, N and sometimes Ca (i.e., in mammals but not insects) are considerably more concentrated (between ∼3.5 - 40×) in animal than in plant biomass (Fig. 2, Table 1). A comprehensive review of taste research reveals that each of these four elements is indeed associated with at least one chemical, or biochemical, known to have a taste that elicits a consumptive response in some animal species. Conversely, we were unable to find evidence of nutritive taste modalities that were obviously associated with specific elements that are closely balanced between plants and animals (except C, see below), or that are more concentrated in plants than in the tissues of the three animal groups in our analysis.

**Fig 2.**
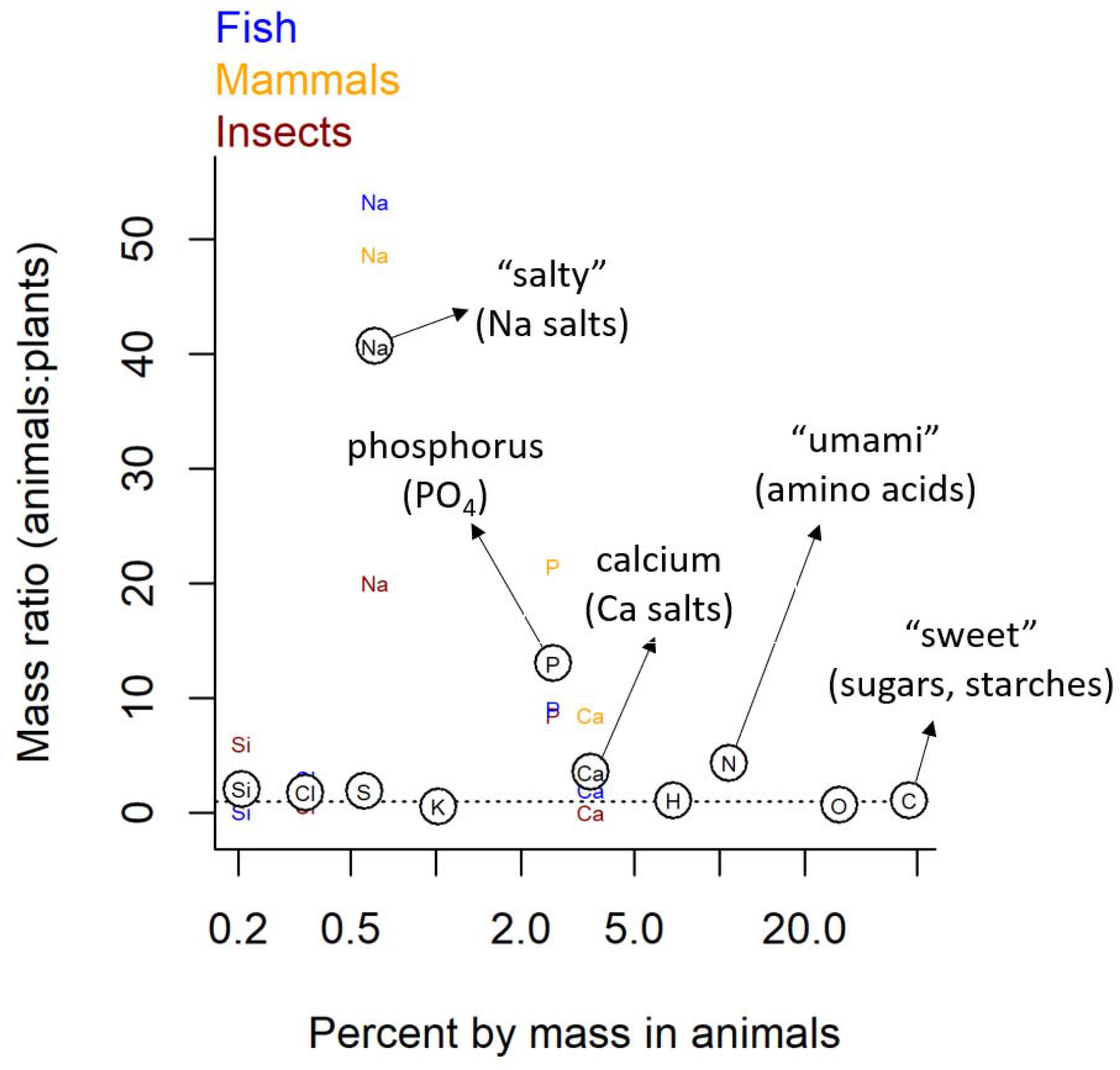
The ratio of biomass concentrations of biologically “essential “ elements in animals and plants for all elements which are at least 0.1% of dry mass in animals. Mass ratio is calculated as: *X*_A_*/ X*_P_; where *X*_A_ and *X*_P_ represent the % of dry mass for element *X* in animals and plants, respectively. Thus, values >1 indicate elements that are more concentrated in animal than plant tissues, 1 indicates equal concentrations and values <1 indicate greater concentrations in plant tissue. Circles containing elemental symbols represent the average of data from mammals (orange), insects (black) and fish (blue) compiled by Bowen (80). Data for plants are from Markert (81) and represent the elemental composition of foliar material.

**Table 1.**
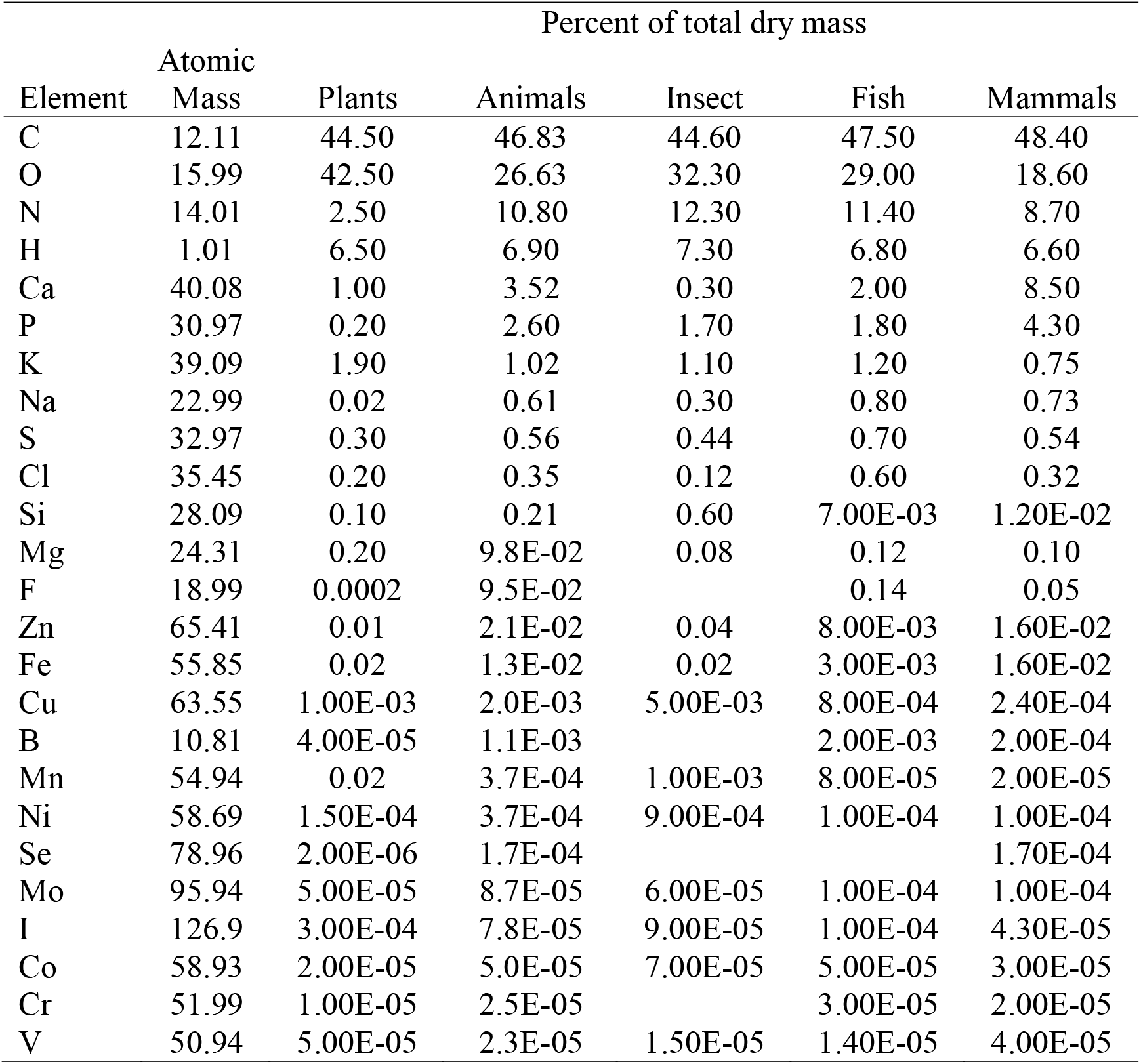
The tissue concentrations (as % of dry mass) of essential elements in plants, animals, insects, fish, and mammals. Values for animals represent the average of elemental concentrations in insects, fish and mammals. Animal data are from Bowen (1980). Plant data are from Markert (81) and are representative of foliar tissue.

For many animals some of these elements (i.e., Ca, Na, P) or associated biochemicals may taste good at some concentrations, but also may produce aversive taste responses when consumed at other, especially high, concentrations. Here again we can turn to the concept of consumer homeostasis to provide a potential explanation for this pattern. Because oversupply of some essential nutrients, such as Na and Ca, is toxic to consumers and because consumers have limited ability to alter their constitution (homeostasis) consumers may be required to regulate homeostasis by managing nutrient intake within a relatively narrow window. The combination of both consumptive and aversive responses to some essential nutrients, especially those that can have toxic or other negative effects when oversupplied, when supplied at different concentrations can be viewed as helping consumers achieve nutritional homeostasis by regulating nutrient intake (3). In the following subsections we discuss taste associations for Na, P, N, and Ca and summarize the evidence that supports these taste capabilities. We provide examples of taste associations, or in some cases preferential feeding behaviors, for these elements within both major vertebrate and invertebrate lineages. Though in some cases direct links between preferential feeding behaviors and associated taste capabilities have not been explicitly made, we believe this anecdotal evidence is at least suggestive of a role for taste in driving these behaviors and warrants further research. We do, however, acknowledge that such behaviors are not necessarily indicative of taste associations, per se, and may be driven by other characteristics of foods such as texture, or by learned nutritional associations (3). Nevertheless, consumptive taste associations for each of these four elements have been described for at least some taxa and we argue that the large number of additional species that display preferential feeding behavior suggests that these taste modalities may be more widespread among animals than what is currently known from research on a relatively limited number of model organisms.

### 3.a. Na imbalances and salty taste

Na comprises ∼0.3% of the body dry mass of animals but only trace amounts of the mass of terrestrial plants, making it on average ∼40× more concentrated in animals than in foliar plant tissue (Fig. 1). Na is typically attractive to both vertebrates and at least some invertebrates at particular concentrations (5, 6, 27, 30, 31, 32). Na salts (i.e., NaCl) are the primary taste molecules associated with gustatory sensing of Na and are responsible for salty taste in humans. The greater Na imbalances between plants and animals than among major animal groups (Fig. 2) suggests that selection for Na taste should be stronger for herbivores than predators (Table 1). However, Na taste remains intact and putatively functional in most mammalian species (12, 13, 33). Nevertheless, herbivorous mammals appear to have a greater affinity for Na than do carnivorous mammals (33), and there is considerable evidence of Na-related feeding behaviors among herbivores (30). For example, many terrestrial herbivores seek out Na-rich mineral deposits on the landscape. Examples include puddling behavior by insects (34), use of dry or wet licks by vertebrates, drinking of sea water by reindeer, soil eating by mountain gorillas (30, 35), and preference for Na rich aquatic plants over terrestrial plants by moose and other ungulates (36). Though Na appears to be attractive to most animals at low to moderate concentrations, aversive responses to high Na concentrations have been reported for both mammals and insects (6, 27, 31). Excess levels of dietary Na salts are detrimental to animals and aversive taste responses to high concentrations can be seen as a mechanism to protect against oversupply. Thus, the evolution of both consumptive and aversive taste responses across Na supply gradients is consistent with biological stoichiometry, as it provides animals sensory cues to guide feeding behavior that maintains relatively tight homeostatic regulation of body Na content, a critical component of osmoregulation. Thus, we suggest that dietary Na imbalances (Fig. 1) combined with the negative consequences of deviation from strict Na homeostasis have driven selection for both consumptive and aversive Na taste responses.

### 3.b. P imbalances and P taste

P imbalances are pervasive at the base of food webs, with P being on average 13× more concentrated in animal than plant biomass (Fig. 2). P availability is often rate-limiting for many organisms and its relationship to growth has garnered considerable interest from a biological stoichiometry perspective (i.e., growth rate hypothesis, 37). Though there has historically been substantial interest among ecologists in the role of P as a limiting element in trophic interactions, taste associations for P-containing compounds have received relatively little attention compared to other basic taste senses (i.e., sweet, salty, bitter, etc.). Nevertheless, P taste has been confirmed in some animal taxa, including cattle (30, 38, 39) and rodents (40), and appears to be elicited most strongly by phosphate, which is the primary form of P in many P-rich biomolecules (i.e., nucleic acids, ATP, phospholipids). Moreover, preferential feeding on P-rich foods has been observed among numerous invertebrate (i.e., grasshoppers, 41) and vertebrate (i.e., rats, 42) taxa and may be fairly widespread. Because generalist herbivores are more likely to suffer greater dietary P deficiencies than carnivores (Fig. 1), and thus stronger selective pressure for P-associated taste, we propose that P taste should be more common among herbivores (and omnivores) than strict carnivores. Additionally, the higher P-requirements imposed by building and maintaining skeletal bone suggest that vertebrates likely experience stronger selection for P taste than do invertebrates. Lastly, excess dietary P supply has been associated with reduced individual growth rates in a variety of organisms (29), reflecting the importance of stoichiometrically balanced P supply for consumers. As such, animals may be likely to find P-associated taste to be aversive at high concentrations, similar to Na and Ca, which may explain why some species reduce consumption rates when reared on high-P diets (crustaceans [43], chickens [44] and mice [40]).

### 3.c. N imbalances and umami taste

N is, on average, ∼4× more concentrated in animal than foliar plant biomass (Fig. 2). The frequently limiting role of N makes it a strong candidate for consumptive taste associations. Indeed, such taste associations for various N-rich amino acids appear to be widespread, having been described in numerous vertebrate and invertebrate taxa. In humans, umami taste is stimulated by the N-rich amino acids L-glutamate and L-aspartate whereas mice are attracted to many L-amino acids (45). Insects also make consumptive responses to a variety of amino acids, although the relative sensitivity to different amino acids varies among insect taxa (46). We posit that selective pressure for N-associated taste should generally be stronger in herbivores than predators, given the greater N-content of animal than plant biomass and thus greater likelihood of N-limitation among herbivores (Fig. 1). However, retention of umami taste in obligate carnivores, may be an indication of the potential for carnivore N-limitation. Despite the appearance of relatively stoichiometrically similar prey, carnivores may experience N-limitation due to their reliance on amino acids as a source of glucose for cellular respiration and the low carbohydrate content of their food (animal prey, 47).

### 3.d. Ca imbalances and Ca taste

Ca is, on average, ∼3.5× more concentrated in the tissues of animals than in plants, although there is major taxonomic variation in the Ca content of animals due to the production of Ca-rich skeletal bone in vertebrates (Fig. 2). Consumptive taste associations for several Ca salts have been observed among diverse groups of animals including amphibians, birds, and mammals (48, 49, 50, 51). Calcium taste is likely to be under stronger selection in herbivores than predators, due to the generally lower Ca content of plant biomass (Fig. 2, Table 1). Moreover, selection for Ca taste should vary considerably between invertebrates and vertebrates due to large differences in Ca-content associated with skeletal bone (and eggshell formation) in the latter. Indeed, Ca appears to be strictly aversive in insects, which are depleted in Ca relative to plants (Fig. 1, [32, 52]), whereas Ca is attractive at low and moderate concentrations to a diverse suite of vertebrates. For example, the Ca content of plant material has been linked to consumptive preferences in rats and mice in laboratory experiments (53), as well as the feeding behaviors of wild gorilla populations, which use forest clearings to access their preferred Ca and Na rich plant species (54). Additionally, dietary Ca deficiencies in herbivorous desert tortoises have been proposed to explain their occasional consumption of numerous Ca-rich resources such as bones, bone-rich vulture feces, reptile skin castings, feathers, mammalian hair (55), and Ca carbonate stones (56). Similar to Na taste, Ca taste is aversive at high concentrations that may result in hypercalcemia (57, 52). Thus, the presence of both consumptive and aversive responses to different levels of dietary Ca supply indicate a role for taste in regulating Ca homeostasis in animals and suggest homeostatic maintenance as a likely driver behind the evolution of Ca-associated taste.

## 4. Carbon, carbohydrate and lipid taste, a nutritional geometry perspective

We have demonstrated that a relatively simple analysis of elemental imbalances between plants and animals, rooted in foundational concepts of biological stoichiometry, explains the context for the evolution of multiple nutritive taste modalities, including those for Na, N (amino acids), Ca and P or associated compounds. However, conspicuously absent from this list are nutritive taste associations for certain C-rich molecules, including the pleasing taste of sugars experienced by humans (sweet taste) and many other organisms. Indeed, biological stoichiometry appears insufficient in providing an explanation for why many animals find certain C-rich biomolecules, such as sugars, pleasing, and others (i.e., structural carbohydrates) not. Carbon is the dominant element by mass in animals and plants and is similarly concentrated in both animal and plant foliar tissue (Fig. 2). However, an organism’s total demand for C is not determined solely by its body C composition but must also account for additional C for energy metabolism (58), given that C compounds are relied upon as energy by nearly all life forms. Thus, focusing strictly on biomass proportions of C in plants and animals suggests that the C concentration of food is unlikely to limit consumer growth and metabolism. As such, dietary C imbalances for consumers may be obscured by the similar C content of animals and plants (Fig. 2).

Assessing potential C-related nutritional imbalances based on bulk C concentration of foods is also difficult given the wide variety of C-containing biochemicals, that vary in their essentiality, digestibility and ability to be synthesized by consumers. For example, a large proportion of the C in foliar tissue of plants occurs in the form of recalcitrant structural carbohydrates (i.e., cellulose and lignin) that many animals cannot digest, rather than structurally simpler carbohydrates, such as sugars and starches, that are more labile and energy dense.

Additionally, many essential C-containing and C-rich macronutrients, including some lipids and amino acids, cannot be synthesized endogenously by animals and therefore must be acquired through the diet. As such, it would be advantageous for animals to differentiate among foods not by their total C-concentration, but by assessing the presence and relative abundances of different C-containing macronutrients.

Here we turn to the framework of nutritional geometry to inform our understanding of how the evolution of C-associated tastes, such as that for sugars, are shaped by the need to regulate nutritional homeostasis (i.e., a target nutrient intake ratio) while foraging among foods that vary in their relative concentrations of multiple essential macronutrients. Nutritional geometry studies have often focused on the relative concentrations of C-containing macronutrients, including carbohydrates, lipids and proteins, in multiple foods. Given that some of these nutrients cannot be synthesized by animals, and that they are variably concentrated in different foods, one would expect that animals would experience strong selective pressure on mechanisms, including taste, that allow them to detect their presence and assess their concentrations in potential foods (3). In the next several paragraphs we discuss the evidence for taste associations for carbohydrates and lipids and how nutritional geometry can inform our understanding of their evolutionary history. Although we previously discussed the evolution of amino acid taste in context of dietary N deficiencies, we acknowledge here that nutritional geometry may provide additional insight into the evolution and radiation of amino acid taste in animals. For example, the more nutritionally explicit framework of nutritional geometry may help us understand why different animal species taste different combinations of amino acid molecules by considering the unique nutrient requirements and diets of different species.

Carbon is the primary energy currency of life and it is glucose that fuels the process of cellular respiration in animals. Dietary carbohydrates, such as sugars and starches, are the primary source of glucose for many animals, though it can also be derived from other sources (i.e., lipids, amino acids). Given that foods can vary substantially in their total carbohydrate content, and that different carbohydrates (i.e., sugars vs. fiber) vary in their digestibility, animals should experience selective pressure on mechanisms that allow them to detect and differentiate between types of dietary carbohydrates in foods. Indeed, many animals taste simple sugars and find sugar taste attractive, while many mammal species are also able to taste, and enjoy, more complex carbohydrates, including starches and starch derivatives (59). Animals, however, do not appear to taste more complex structural carbohydrates, such as cellulose fiber. Thus, it appears that carbohydrate taste is fine-tuned for labile, energy-rich carbohydrates, thereby enabling consumers to regulate their intake of energetically important foods. The link between sugar taste in animals and metabolic C demand that we suggest is supported by multiple lines of evidence. For example, in frugivorous non-human primates, sucrose taste threshold increases with decreasing body size (60), indicating that the sweet taste receptors of small-bodied species are tuned for selecting foods with higher sucrose than those of large-bodied species. This pattern is consistent with basic allometric scaling rules relating to organismal metabolic rates, where mass-specific metabolic rates decrease with increasing body size (61) and is suggestive of a link between sugar taste thresholds and C demands of at least some primate species. Moreover, bumble bees and honey bees have gustatory receptor neurons that are triggered by especially high sugar concentrations (62) providing a gustatory mechanism to meet their high metabolic C demand. Likewise, hummingbird’s high metabolic demands for C are consistent with the evolution of their sweet taste receptor from their ancestor’s umami taste receptor as their diet and habits changed (63). These patterns are consistent with the principles of nutritional geometry which recognizes animals use sensory systems, like taste, to inform foraging decisions that balance the intake of multiple nutrients that vary in their concentration among different foods.

Lipids, including fatty acids, phospholipids and steroids, are another important and diverse class of C-rich macromolecules that are essential in animal nutrition and can be an important source of glucose for cellular respiration. Potential foods (i.e., nuts and seeds vs fruits and foliage, plant vs animal prey) may vary substantially in the both the amount and types of lipids that they contain and some nutritionally important lipids, including various essential fatty acids, must be obtained directly from the diet. As such, animals should benefit from the ability to evaluate the lipid composition of different foods to maintain a balanced intake of essential lipids and other nutrients relative to nutritional demands. Indeed, animals possess multiple systems, including taste, olfaction, and gut-nutrient sensing, that contribute to lipid detection in their foods and lipids are increasingly recognized as having pleasing “flavor”, in addition to smell and texture, for many animals (64). However, evidence for specific positive taste associations for major lipid molecules, such as various fatty acids, is not widespread among animals, indicating that other mechanisms such as olfaction and gut nutrient sensing may be more important for detecting dietary lipids and driving lipid intake in some species. For example, taste for fatty acids appears to be aversive in many species, including humans, and the attractive taste of lipids experienced by other species, including mice, is apparently stimulated only by certain long chain fatty acid molecules, such as linoleic acid (64, 65), an essential polyunsaturated omega-6 fatty acid that must be acquired through the diet. Thus, fatty acid taste in some species may be seen primarily as a mechanism for regulating consumption of specific essential fatty acids, which animals cannot synthesize endogenously, a pattern that is consistent with concepts from nutritional geometry. The pleasing sensation experienced when consuming dietary fats by some species may be triggered primarily by sensory cues other than taste [i.e., olfaction, texture, postingestive cues (66)], thereby highlighting the need to consider how other sensory systems interact to influence foraging behaviors that play a role in regulating the intake of lipids (3).

Nutritional geometry implies that animals that consume a variety of foods of varying nutrient composition will benefit from the ability to detect, whether by taste or another mechanism, the presence of multiple nutrients in their food. Animals should experience particularly strong pressure for nutrient sensing systems that detect essential biochemicals that cannot be synthesized and must be consumed. Tastes for certain fatty acids, as discussed above, are examples. Another example is the attractive taste of vitamin C (ascorbic acid) to guinea pigs (67) and certain laboratory strains of rats (68), which are incapable of vitamin C synthesis. Consumptive taste associations for vitamin C, which has a chemical formula of C_6_H_8_O_6_, would not necessarily be predicted from an analysis of stoichiometric imbalances between guinea pigs or rats and their food.

## 5. Lack and loss of taste senses among animal species

Understanding why some animal species possess, or lack, certain taste senses has generated considerable interest among taste researchers, particularly in cases where specific taste functions have been lost within some animal lineages (12, 13). The loss of taste receptor function has been linked, in part, to evolutionary shifts in feeding ecology, including dietary specialization, that reduce the selective pressures on taste receptor genes (11, 12, 69, 70, 71, 72). For example, sweet taste has been independently lost in multiple lineages of terrestrial carnivores, including the strictly carnivorous felids, cetaceans, and vampire bats (12, 13, 71). This loss of sweet taste has been linked to pseudogenization of the T1R2 (sweet) receptor gene following dietary shifts towards obligate carnivory (13) as the relative lack of carbohydrates in animal tissue is thought to reduce selective pressure for maintaining a functioning sweet taste receptor (12). Shifts in feeding mode, meaning how an organism feeds (i.e., swallowing versus chewing prey, fluid feeding, etc.), have also been linked to loss of taste functions in some animal lineages. For example, the substantial loss of taste (sweet, umami, bitter and sour) in cetaceans has been linked to pseudogenization of multiple genes following an ancestral dietary shift from herbivory to carnivory (animals contain fewer carbohydrates and bitter tasting compounds than plants), as well as a behavioral shift towards swallowing prey whole, which precludes the release of taste compounds within the oral cavity through mastication (72). Additionally, in vampire bats, the shift to blood feeding and use of infrared sensing to locate blood flows have presumably reduced selection for functional carbohydrate and amino acid taste receptors (71) resulting in the loss of sweet and umami tastes, respectively. Despite these notable examples, lineage-specific pseudogenization events do not always align with shifts in feeding ecology or behavior, suggesting gaps in our understanding of the physiological function of taste receptor genes and the selective pressures that influence their maintenance (13, 73).

We suggest that biological stoichiometry can provide additional insight into the ecological contexts under which natural selection is likely to favor, or possibly disfavor, the evolution and maintenance of functional taste capabilities across diverse animal lineages. By focusing on differences in the elemental composition of plants and animals, as well as major animal lineages, we can use biological stoichiometry to understand how selective pressure for certain taste capabilities might vary among species on the basis of their trophic position (herbivore, omnivore, or carnivore) and phylogenetic relationships (i.e., mammals and insects). For example, differences in organismal stoichiometry among trophic levels, such as those depicted in Fig. 1, suggest that animals are likely to experience differential selective pressure for certain taste capabilities based on their trophic position. The narrower variation in the elemental composition of animals compared with plants (Fig. 1) indicates that stoichiometric imbalances for N and P, and potentially Na and Ca (Fig. 2), are likely to be greater, and more common, for animals that feed at lower trophic positions, such as generalist herbivores and omnivores, than those that feed at higher trophic levels (i.e., obligate carnivores). Put another way, carnivores are more likely to encounter food that has a chemical composition matching their own, thereby reducing selective pressure on the genes that allow them to differentiate among foods of varying composition. Conversely, generalist herbivores and omnivores consume foods that are much more heterogeneous in their elemental composition, and less likely to be in balance with their own chemical composition, meaning they should experience stronger selective pressure on their gustatory systems than strict carnivores.

When taste receptor genes are no longer under strong selective pressure, they should be more likely to accumulate deleterious mutations that result in their becoming broken pseudogenes. To illustrate this concept, we turn to data compiled by Feng and Zhao (73) who used available genomes of 48 mammals to demonstrate widespread pseudogenization of taste receptor genes, including those for the T1R1 umami receptor. Based on principles of biological stoichiometry, we would expect herbivores to experience much greater dietary N-deficiencies than carnivores, because of the higher and more variable C:N ratio of plants than animals (Fig. 1), and therefore stronger selective pressure to maintain functioning umami (amino acid) taste receptor genes. Though Feng and Zhao (73) concluded that there was no common dietary factor that explained the pseudogenization of the T1R1 umami taste receptor across those 48 mammal species, it is true that a higher proportion of predatory species (carnivores, insectivores, piscivores) lacked a functioning umami receptor gene (60% defective or absent, n=10) than omnivores (13.5% defective or absent, n=22), or herbivores (31% defective or absent, n=16) in their dataset. Thus, there is some limited evidence that N-associated umami taste is more common for consumers that experience large dietary imbalances in N-supply, suggesting that biological stoichiometry may be useful for understanding general patterns regarding the loss of some taste functions in animals. However, we caution that biological stoichiometry may not accurately predict the presence or absence of specific tastes for individual species as those are undoubtedly shaped by each species unique evolutionary history. For example, the loss of amino acid taste in some herbivores, like the giant panda, would not be expected when viewed from the perspective of biological stoichiometry. In the case of the giant panda, loss of umami taste is likely a reflection of reduced selective pressure on taste receptors resulting from the reduced dietary breadth of giant pandas, which almost exclusively consume bamboo, relative to ancestral species (11). Additionally, retention of umami taste in some carnivores, such as felids, may be an indication of predator N-limitation, despite the appearance of relatively stoichiometrically balanced prey (47, 74). In such cases, N-limitation may be driven by consumer metabolism, as many predators rely on gluconeogenesis to convert N-rich amino acids into glucose for cellular respiration due to the lack of carbohydrates in animal prey (47).

Though interspecific variation in elemental composition tends to be lower among different animal than plant species, there are several elements that vary considerably among animal species based on broad scale phylogenetic relationships. For example, P and Ca tend to be more concentrated in vertebrates than insects, a result of the presence P-and Ca-rich skeletal bones in the former (Fig. 1). As such, vertebrates may be more likely to experience, or experience greater magnitude, P and Ca imbalances than insects within any given trophic level, making it more likely they should find P and Ca taste attractive. Indeed, Ca is attractive at some concentrations to a diverse suite of vertebrates (birds, amphibians, mammals), but appears to be strictly aversive in insects, which are depleted in Ca relative to plants (Fig. 1; 32, 52). Thus, we believe that biological stoichiometry can inform our understanding of the phylogenetic distribution of taste by revealing differential elemental imbalances between animals and their food both among major animal lineages as well as consumer trophic levels, while accepting that the full suite of taste capabilities possessed by any species will ultimately be influenced by that species’ evolutionary history.

## 6. Limitations of biological stoichiometry and nutritional geometry

We do not intend to provide a comprehensive review of taste biology research, as several such papers exist and remain useful (i.e., 5, 6), nor do we intend to provide a comprehensive synthesis of either biological stoichiometry theory or nutritional geometry. Rather, our intent here is to highlight linkages between taste and animal nutrition using two prominent nutritional frameworks in ecology. We do, however, draw support for this effort from a comprehensive review of the taste literature, which in some cases (i.e., for P- and Ca-related taste) is limited to a relatively small number of species and model systems, including humans, mice and livestock. As such, some of the evidence we present to support our general thesis, that biological stoichiometry and nutritional geometry can inform our understanding of the evolutionary contexts that produced multiple nutritive taste modalities, is anecdotal in nature. We believe that these examples suggest a possible role for taste in driving the documented behaviors, especially in species for which specific taste capabilities have not been previously investigated. It is important that we acknowledge that animals employ other sensory systems, as well as learned nutritional associations, to assess the nutritional content of their foods and that these systems may also contribute to selective feeding behaviors (3, 66). Finally, we acknowledge that both biological stoichiometry and nutritional geometry may not accurately predict the full suite of consumptive taste capabilities for a given species, but nonetheless are useful for generating hypotheses to further understand taste evolution.

We have previously discussed that dietary specialization has been linked to reductions in diversity of taste capabilities in some species. In such cases, the presence or absence of taste senses may deviate from expectations based solely on principles of biological stoichiometry and nutritional geometry. In cases of extreme dietary specialization, especially those that result in large nutritional imbalances between the consumer and its food, animals may rely on a diverse suite of adaptations ranging from specialized physiology and digestive morphology, to mutualisms with endosymbiotic microorganisms to reduce the severity of nutritional deficiencies in their diets. For example, some termites that feed on nutritionally poor wood rely on microbial endosymbionts to breakdown cellulose (75), and for microbial N-fixation within the gut (76). Animals that possess such adaptations may circumvent the effects of nutritional limitation on poor quality diets by means other than dietary selectivity informed by gustatory sensing, and therefore may experience little selective pressure to maintain functioning gustatory systems.

We have primarily focused our discussion of taste senses on those that promote consumptive, rather than aversive, behavioral responses, though we do discuss aversive responses to high concentrations of compounds that produce consumptive responses under some conditions (i.e., Na, P and Ca salts). This distinction is important given that some aversive tastes, such as bitter and, perhaps to some extent sour, are generally thought to inform consumers that foods may contain toxic compounds, or low pH, that may be harmful if ingested, even in low quantities (5, 6). Thus, aversive tastes encourage consumers to reject foods, and do not necessarily align with our applications of biological stoichiometry theory and nutritional geometry, which we link primarily to consumptive tastes that drive nutrient acquisition. Indeed, aversive tastes may alert consumers to the presence of potentially harmful compounds in foods that may appear nutritious when considered from a strictly biological stoichiometry perspective. Interestingly, animals can, in some cases, learn to tolerate, or even prefer, bitter and sour tastes in their food and drink, particularly when the toxicity of taste compounds is negligible relative to potential nutritional and pharmacological benefits (77). Though aversive responses to bitter and sour tastes are not necessarily consistent with principles of biological stoichiometry, we have documented several examples where aversive tastes for high concentrations of certain elements, such as Na, P and Ca, do align. The aversive responses to highly concentrated supply of Na, P and Ca contrast to the consumptive response that these same elements induce at low and moderate concentrations. These differential responses to low and high levels of dietary Na, P and Ca can be viewed as adaptations that reduce the potential for physiological consequences associated with deviation from strict homeostatic Na, P and Ca regulation, and thus provide examples of aversive taste responses that are consistent with the core principles of biological stoichiometry theory.

## 7. Future directions

We have provided numerous hypotheses for understanding the evolution of consumptive taste in animals based on the core principles of biological stoichiometry theory and the complementary framework of nutritional geometry. We believe that our review of the taste biology literature, albeit brief and incomplete, has produced compelling evidence that supports the usefulness of these frameworks for informing our understanding of the evolution of taste. Here, we highlight several observations and questions that have emerged from this analysis that point to exciting avenues for future work that we hope will bridge the fields of taste biology, physiology, biological stoichiometry, ecology, and evolution.

We have previously acknowledged that support for some of the ideas presented in this manuscript come from relatively small bodies of literature. For example, both P and Ca-associated tastes have been described for a small number of taxa, reflecting the lack of attention for these particular taste capabilities relative to the basic taste modalities recognized in humans (e.g., salty, bitter, sour, sweet, umami). Nevertheless, our analysis suggests that P- and Ca-associated taste may be widespread given the large number of species that display preferential feeding behaviors related to these elements, and provides some guidance on where to look for candidate species that might possess these specific taste capabilities (i.e., herbivores and omnivores for P, vertebrates for Ca). Thus, we believe that both P- and Ca-associated tastes are deserving of increased attention in a wider array of taxa, including invertebrates.

In addition to the elements surveyed here (Na, P, N, Ca and C), several trace elements such as Bo, Fl, Fe and Zn are present in much greater concentrations in animal tissues than the foliar tissue of plants, suggesting the potential for consumptive taste associations for those elements. For example, it is worth noting that Zn is essential for the proper functioning of taste receptors themselves (78). Future research may assess the potential for gustatory sensing of those elements in animals, followed by efforts to characterize the genetic and molecular nature of those gustatory systems. Further research is also warranted to better understand potential taste associations for essential biochemicals that might not be predicted by biological stoichiometry. Gustatory sensing of essential biochemicals, particularly those that are not endogenously synthesized by animals, should be highly favored leading to the evolution of taste associations.

Rapid advances in molecular and genetic techniques have enabled great insight into the chemical and genetic underpinnings of taste in both vertebrate and invertebrate models. We now know the specific receptor genes involved in producing many different taste modalities. This opens the possibility of analyzing the distribution of different taste modalities throughout the animal kingdom and thereby facilitating the extension of taste research to a larger number of species, including a greater survey of invertebrate fauna. This might prove especially useful for ecologists seeking to understand how animals employ behavioral or life history strategies to minimize nutritional imbalances in natural settings, thereby influencing interspecific predator-prey interactions, food web and ecosystem dynamics. For example, a species ability to taste a particular nutrient is likely an important trait that influences the role of that species in nutrient cycling. We encourage continued efforts to understand the genetic and molecular bases of different taste capabilities, particularly those exhibited by invertebrates. Identification of specific taste receptors and their associated genes will facilitate broad phylogenetic analyses to reveal the taxonomic distribution and evolutionary history of individual taste capabilities.

## Data Accessibility

Data sharing not applicable to this article as no datasets were generated and all data presented (Table 1, Fig.1, Fig.2) can be accessed from the original sources.

## Author contributions

LMD, BWT, BJR and RRD were primarily responsible for the conceptual development of the manuscript and MGT provided guidance and expertise on discussion of taste biology. LMD compiled relevant data and created figures. All authors made writing contributions during manuscript preparation and were involved in editing and revision of various drafts.

## Acknowledgements

This manuscript was improved by comments from P Jiang, JA Balik, R Irwin, and S Jordt. This work was supported by the U.S. National Science Foundation [grant number 1556914].

